# Epstein-Barr virus nuclear antigen 2 (EBNA2) extensively rewires the human chromatin landscape at autoimmune risk loci

**DOI:** 10.1101/2020.04.15.043612

**Authors:** Ted Hong, Sreeja Parameswaran, Omer Donmez, Daniel Miller, Carmy Forney, Michael Lape, Mariana Saint Just Ribeiro, Jun Liang, Lee E. Edsall, Albert Magnusen, William Miller, Iouri Chepelev, John B. Harley, Bo Zhao, Leah C. Kottyan, Matthew T. Weirauch

## Abstract

The interplay between environmental and genetic factors plays a key role in the development of many autoimmune diseases. In particular, the Epstein-Barr virus (EBV) is an established contributor to multiple sclerosis, lupus, and other disorders. Previously, we demonstrated that the EBV nuclear antigen 2 (EBNA2) transactivating protein occupies up to half of the risk loci for a set of seven autoimmune disorders. To further examine the mechanistic roles played by EBNA2 at these loci on a genome-wide scale, we globally examined gene expression, chromatin accessibility, chromatin looping, and EBNA2 binding, in a B cell line that was 1) uninfected, 2) infected with a strain of EBV lacking EBNA2, or 3) infected with a strain that expresses EBNA2. We identified >400 EBNA2-dependent differentially expressed human genes and >5,000 EBNA2 binding events in the human genome. ATAC-seq analysis revealed >2,000 regions in the human genome with EBNA2-dependent chromatin accessibility, and HiChIP-seq data revealed >1,700 regions where EBNA2 altered chromatin looping interactions. Importantly, autoimmune genetic risk loci were highly enriched at the sites of these EBNA2-dependent chromatin-altering events. We present examples of autoimmune risk genotype-dependent EBNA2 events, nominating genetic risk mechanisms for autoimmune risk loci such as *ZMIZ1*. Taken together, our results reveal important interactions between host genetic variation and EBNA2-driven disease mechanisms. Further, our study highlights a critical role for EBNA2 in rewiring human gene regulatory programs through rearrangement of the chromatin landscape and nominates these interactions as components of genetic mechanisms that influence the risk of multiple autoimmune diseases.

## Introduction

Crosstalk between genetic risk polymorphisms and environmental factors is thought to influence the onset and progression of many human diseases (Hunter 2005; Bookman et al. 2011; McAllister et al. 2017). Many diseases have a complex genetic etiology, including cancers (Flavahan et al. 2017), cardiovascular diseases (North et al. 2003), and autoimmune diseases such as multiple sclerosis (MS) (Ascherio and Munger 2007) and systemic lupus erythematosus (SLE) (Kamen 2014). Over the past 15 years, a multitude of genome-wide association studies (GWASs) have identified more than 50,000 disease-associated genetic variants for many disorders (Tam et al. 2019). As many as 90% of disease-associated genetic variants fall within non-coding regions of the genome (Hindorff et al. 2009; Maurano et al. 2012), implicating a key role for regulatory proteins such as transcription factors (TFs) in the etiology of human disease (Lee and Young 2013; Deplancke et al. 2016). Regulatory proteins bind to promoter regions and distal regions of target genes (e.g., enhancers) to alter gene expression through numerous mechanisms (reviewed in (Lambert et al. 2018; Sullivan et al. 2018; Schoenfelder and Fraser 2019)). Some regulatory proteins, such as the pioneer factor EBF1, are capable of directly altering the chromatin landscape. Other TFs, such as YY1 and CTCF, can affect the three-dimensional structure of chromatin by facilitating the formation of novel chromatin loops that alter gene transcription (Beagan et al. 2017; Weintraub et al. 2017). Thus, regulatory proteins likely can contribute to human disease processes through a variety of mechanisms.

Viral infections are a common environmental exposure known to be closely linked to many human diseases (Bray et al. 1983; Foxman and Iwasaki 2011; Hong et al. 2014; Pender and Burrows 2014). In particular, previous studies have revealed causative roles for the Epstein-Barr virus (EBV) in mononucleosis (Dunmire et al. 2015), Burkitt’s lymphoma (Rochford and Moormann 2015), and Hodgkin lymphoma (Vockerodt et al. 2014). EBV is also strongly implicated in autoimmune diseases such as rheumatoid arthritis (RA) (Balandraud and Roudier 2018), inflammatory bowel disease (IBD) (Dimitroulia et al. 2013), SLE (Harley and James 2006), and MS (Bagert 2009). Despite extensive epidemiologic and serological evidence, the molecular mechanisms through which EBV-host interactions increase autoimmune disease risk remain largely unknown.

Viruses can directly perturb the host’s transcriptome through the actions of virus-encoded transcriptional regulatory proteins (Agudelo-Romero et al. 2008; Clyde and Glaunsinger 2010; Bermudez-Morales et al. 2011; Graham 2016; Harley et al. 2018; Liu et al. 2020). Viral transcriptional regulators can either interact with the host genome directly, as is the case for the EBV-encoded Zta protein (Flemington and Speck 1990; Mahot et al. 2003), or indirectly through interactions with host DNA-binding factors, such as the EBV-encoded Epstein-Barr virus nuclear antigen 2 (EBNA2) protein and the human TF RBPJ (Henkel et al. 1994). In both cases, genetic variation in the host genome can affect these virus-host interactions, leading to alteration of host gene expression levels (Bochkov et al. 2010; Caliskan et al. 2015; Harley et al. 2018).

EBNA2 controls multiple processes, including the immortalization of EBV-infected B cells, by altering the expression levels of human genes (Pich et al. 2019). Mechanistically, EBNA2 mediates at least some of this regulation through interactions with human TFs such as RBPJ, SPI1 (PU.1), and EBF1 (Zhao et al. 2011). The EBF1 protein can bind to and open chromatin that is occupied by EBNA2-RBPJ complexes (Lu et al. 2016). EBNA2 can also recruit chromatin remodelers such as histone acetyltransferase p300/CBP (Wang et al. 2000; Lu et al. 2016) and the SWI/SNF complex (Wu et al. 2000), further supporting a role for EBNA2 in human chromatin rearrangement. Likewise, the EBNA2-RBPJ complex can create a new chromatin looping interaction between a distal enhancer region and the *MYC* promoter, inducing *MYC* expression that leads to continuous B cell proliferation (Zhao et al. 2011; Wood et al. 2016; Jiang et al. 2017). Despite these strong independent lines of evidence implicating EBNA2 in the alteration of chromatin accessibility and looping in the human genome, a genome-wide investigation of EBNA2-dependent human chromatin alteration has not been previously performed.

A recent study from our group revealed that a significant number of autoimmune disease-associated genetic loci contain genetic variants that are located within EBNA2 ChIP-seq peaks (Harley et al. 2018). In particular, nearly half of the SLE and MS genetic risk loci contain disease-associated genetic variants that are directly located within regions of the human genome occupied by EBNA2. We also discovered numerous examples of autoimmune-associated genetic variants that alter the binding of EBNA2 and other transcriptional regulators to the human genome in a genotype-dependent manner. Collectively, these results are consistent with EBNA2 playing an important role in autoimmune disease etiology.

Understanding the molecular mechanisms mediating interactions between EBNA2 and the human genome is important for achieving an understanding of the development and progression of autoimmune diseases. In our previous study, analysis of public datasets revealed the presence of EBAN2 ChIP-seq peaks at up to half of the risk loci for particular autoimmune diseases. In this study, we explore the role of EBNA2 within the human B cell gene regulatory network by examining EBNA2-dependent alterations to the human chromatin landscape and investigating the impact of autoimmune disease-associated genetic polymorphisms on these mechanisms. To identify EBNA2-dependent effects, we employ an experimental design comparing human B cells that are either 1) uninfected, 2) infected with an EBV strain that lacks EBNA2 (EBV^EBNA2−^), or 3) infected with an EBV strain that has EBNA2 (EBV^EBNA2+^). Using this approach, we identify human genes whose expression level changes coincides with the presence EBNA2 (RNA-seq), and we resolve the effects of EBNA2 on chromatin accessibility (Assay for Transposase-Accessible Chromatin sequencing or ATAC-seq) and chromatin looping (protein-centric chromatin conformation sequencing or HiChIP-seq). Further, we examine the enrichment of autoimmune disease risk genetic variants within these EBNA2-dependent regulatory mechanisms, and identify allele-dependent EBNA2 behavior at autoimmune-associated variants.

## Results

### EBNA2 modulates human gene expression in EBV-infected B cells

To globally measure the effect of EBNA2 on human gene expression patterns, we performed RNA-seq in Ramos B cells in three experimental conditions - uninfected, EBV^EBNA2−^, and EBV^EBNA2+^ (Fig. 1). We used the P3HR-1 EBV strain, which contains a naturally occurring EBNA2 deletion, for EBNA2-infections, and the widely-used B95.8 EBV strain for EBNA2+ infections. We chose to use the immortalized Ramos B cell line instead of primary B lymphocytes as host for the infections to control for the heterogeneity of the B cell compartment. We first examined the presence and absence of EBV-encoded molecules such as EBNA2 and the Epstein-Barr virus-encoded small RNAs (EBERs) across the three cell types. As expected, we only detected transcripts for EBERs in the EBV infected Ramos cells (both EBV^EBNA2+^ and EBV^EBNA2−^), and we only detected EBNA2 transcripts and protein in EBV^EBNA2+^ cells (Supplemental Fig. S1). Further, no RNA-seq reads mapped to the EBV genome in the uninfected dataset, whereas 7535/7764 and 4249/4423 reads mapped for EBV^EBNA2+^ replicate 1/replicate 2 and EBV^EBNA2−^ replicate 1/replicate 2, respectively.

**Figure. 1.**
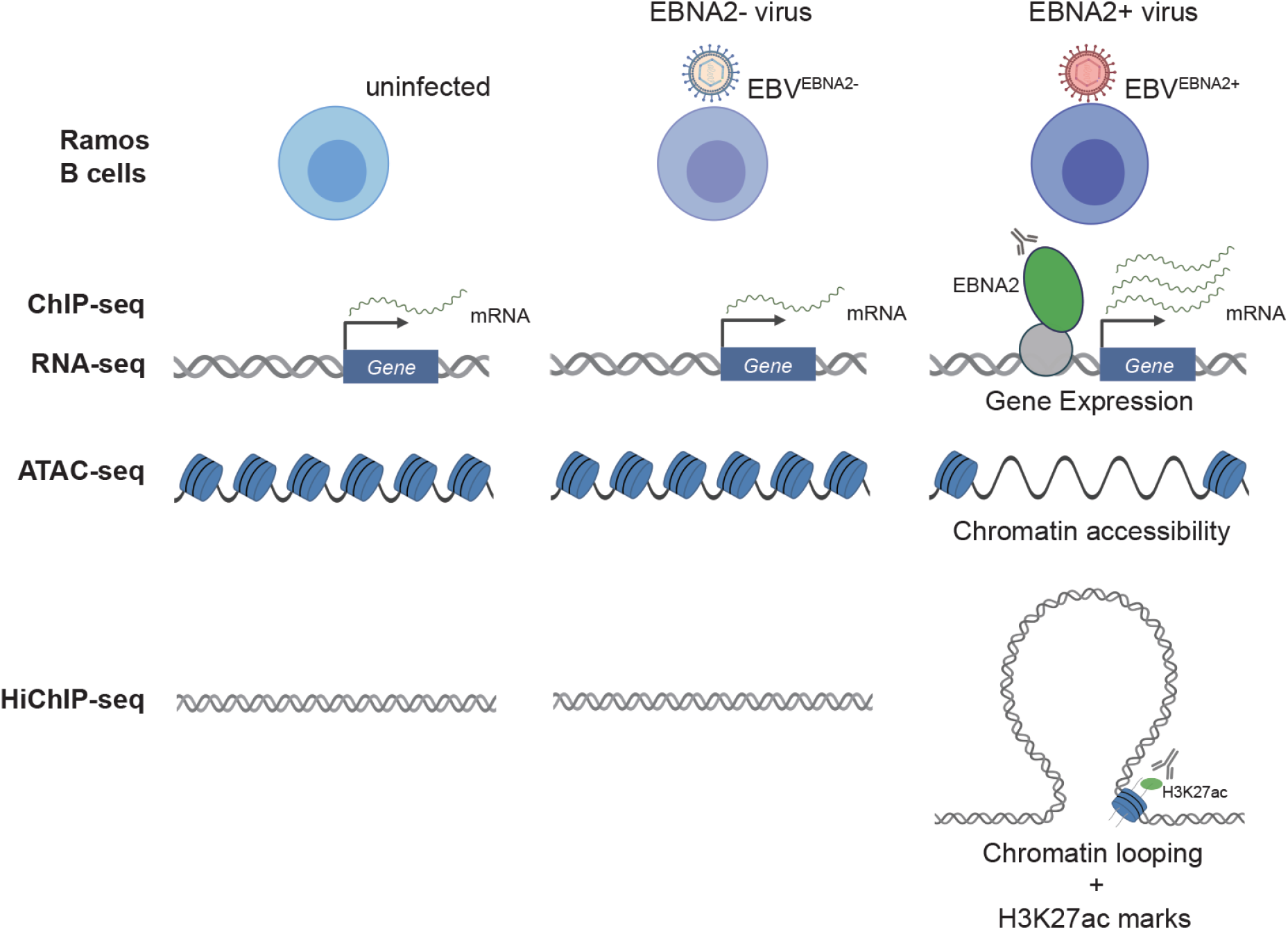
Schematic overview of the experimental design. Our working hypothesis is that EBNA2 alters human gene expression by rewiring the chromatin landscape. To test this hypothesis, RNA-seq, ChIP-seq, ATAC-seq, and HiChIP-seq experiments were performed in uninfected, EBV^EBNA2−^-infected, and EBV^EBNA2+^-infected Ramos B cells.

To investigate the effect of EBNA2 on host gene expression, we identified differentially expressed genes (DEGs) using these three experimental conditions. First, we compared gene expression changes between EBV^EBNA2+^ and uninfected, which captures the effect of EBV infection on human gene expression in B cells. In total, 493 human genes were differentially expressed upon EBV^EBNA2+^ infection (Supplemental Table S1), with 290 genes up-regulated and 203 down-regulated (1.5-fold change or more, adjusted *p*-value < 0.05). Among these EBV-dependent differentially expressed genes, 67 of the 290 upregulated genes and 18 of the 203 downregulated genes were consistent with a previous study examining EBV infection in primary B cells at day 28 post-infection (Wang et al. 2019) (*p*-value: 0.0208, Fishers Exact Test) (Supplemental Table S1).

Next, we identified the EBNA2-specific effects of these gene expression changes. To this end, we identified significant changes in gene expression (1.5-fold change or more, adjusted *p*-value < 0.05) between EBV^EBNA2+^ and uninfected cells, and between EBV^EBNA2−^ and uninfected cells (Figs. 2A and 2B, Supplemental Fig. S2A and S2B, Supplemental Table S1). This procedure identified 421 genes that are differentially expressed in the EBV^EBNA2+^ condition but not in the EBV^EBNA2−^ condition (243 up-regulated genes and 178 down-regulated genes, Supplemental Table S1), which we designate the EBNA2 DEGs. As expected, GO Biological Process enrichment analysis for EBNA2 DEGs revealed processes involved in the immune response, including response to virus, lymphocyte activation, and cytokine production (Supplemental Fig. S3, Supplemental Table S2). Further, a significant proportion of the EBNA2 DEGs, including *CD80, MAP3K8, SLAMF1*, and *ZMIZ1*, are involved in leukocyte cell-cell adhesion (adjusted *p*-value: 6.03E^−3^, Supplemental Table S2). Collectively, these results indicate that the expression levels of hundreds of human genes are affected by the presence of EBNA2.

**Figure 2.**
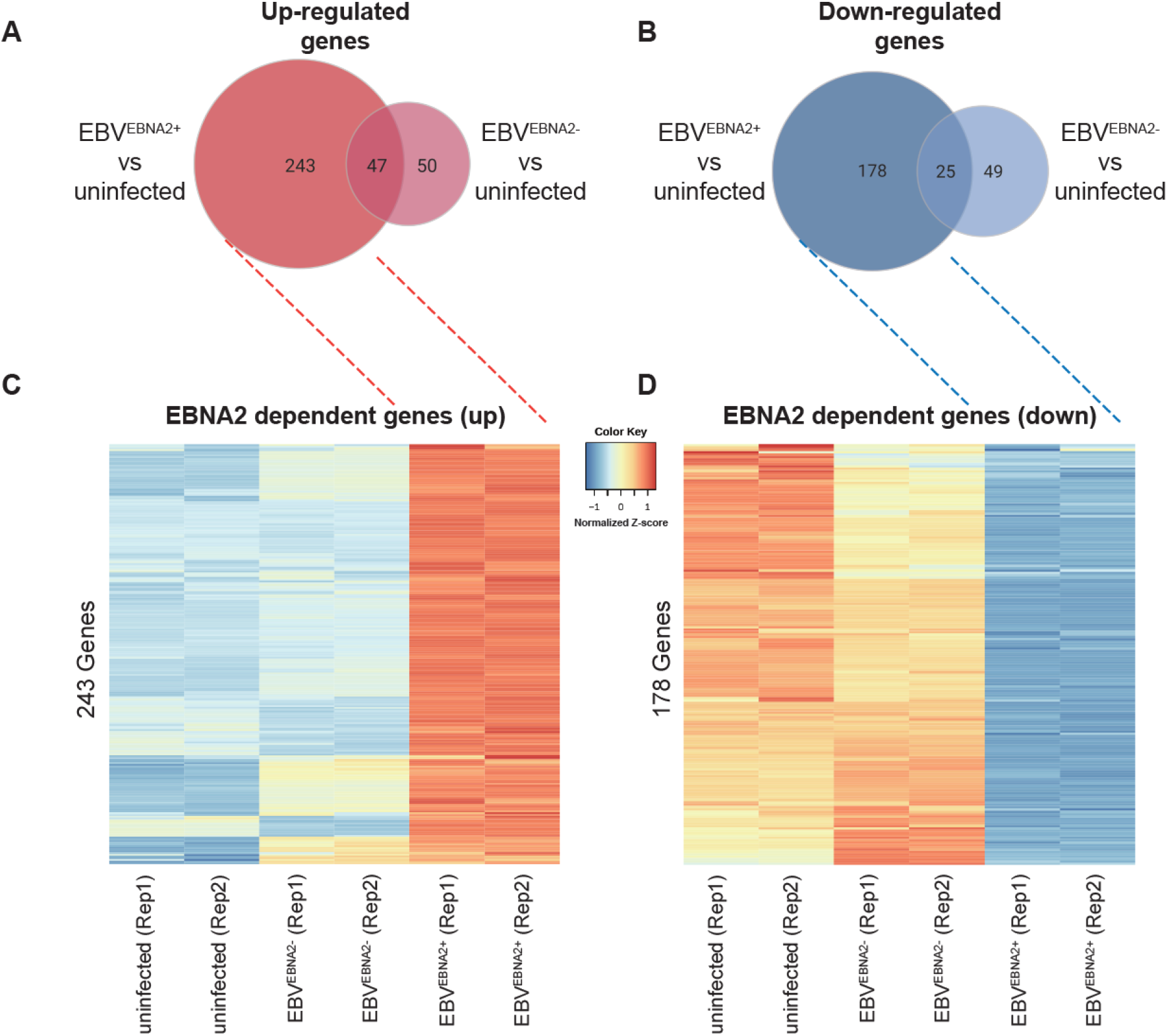
Differential gene expression in EBV-infected Ramos cells. Venn diagrams depicting the number of up-regulated genes (A) and down-regulated genes (B) based on comparisons between EBV^EBNA2+^ vs uninfected and EBV^EBNA2−^ vs uninfected conditions, respectively. Heatmaps depict genes that are specifically expressed (higher or lower) in EBV^EBNA2+^ cells. Values in the heatmaps indicate the normalized relative Z-score of the FPKMs across each row (i.e., the default normalization method in the R ‘heatmap’ function).

### EBNA2 occupies regions of the human genome proximal to genes with EBNA2-dependent expression levels

We next performed ChIP-seq for EBNA2 in EBV^EBNA2+^ Ramos cells, identifying 5,781 regions of the genome occupied by EBNA2 (see Methods). We also performed ChIP-seq for EBNA2 in GM12878 cells, an EBV-transformed lymphoblastoid cell line. Quality control analyses indicated high data quality (Supplemental Table S3) and strong agreement between experimental replicates (Supplemental Fig. S4). Throughout this study, we use our RELI tool (Harley et al. 2018) to compare genomic datasets. In brief, RELI uses a simulation-based procedure to systematically gauge the significance of the intersection between a set of input genomic regions (e.g., EBNA2 ChIP-seq peaks) and each member of a large library of functional genomics experiments (e.g., published ChIP-seq or ATAC-seq peaks). Comparison of the genomic coordinates of our EBNA2 ChIP-seq datasets to published EBNA2 ChIP-seq datasets using our RELI algorithm revealed highly significant concordance (Supplemental Table S4). Likewise, our EBV^EBNA2+^ and GM12878 ChIP-seq peaks aligned significantly with published ChIP-seq peaks of established EBNA2 partners and co-regulators performed in EBV-infected B cells (Zhou et al. 2015), including RBPJ, NFKB1, and EBF1 (Supplemental Table S4). As expected, we also observed enrichment within our EBNA2 peaks for the DNA binding motifs of established EBNA2 partners, such as RBPJ, EBF1, and SPI1 (PU.1) (Supplemental Table S5). EBNA2 peaks are strongly enriched within 100kb, 10kb, and 5k of EBNA2-dependent gene transcription start sites (Supplemental Table S6). Collectively, these results indicate that our EBNA2 ChIP-seq experiments are of high quality.

Next, we investigated the relationship between EBNA2 binding and gene expression changes in Ramos cells using RELI. As expected, EBV^EBNA2+^ Ramos ChIP-seq peaks were enriched within both proximal (promoter, up to 5kb from the transcription start site: 2.7-fold enrichment, adjusted *p*-value: 1.13E^−8^) and distal (enhancer, up to 100kb from the transcription start site: 1.5-fold enrichment, adjusted *p*-value: 3.35E^−8^) regions of EBNA2 DEGs. EBNA2 ChIP-seq peaks were more enriched near up-regulated than down-regulated genes within promoters (3.2-fold and 2.0-fold, respectively) and distal regions (1.7-fold and 1.2-fold, respectively) (Supplemental Table S6). For example, EBNA2 ChIP-seq peaks are located in the promoter region of EBNA2 DEG *LY9* (Supplemental Fig. S5), in agreement with a recently published finding that *LY9* gene expression is induced by EBV infection (Wang et al. 2019). LY9 protein expression levels have also been shown to be increased in the presence of SLE immune complexes (Hagberg et al. 2013), suggesting a possible role for EBNA2 in *LY9* gene regulation in lupus.

### EBNA2 alters chromatin accessibility at hundreds of human genomic loci

We next used our Ramos experimental system to examine the impact of EBNA2 on genome-wide chromatin accessibility by performing ATAC-seq in uninfected, EBV^EBNA2−^, and EBV^EBNA2+^ conditions (see Methods). In total, we identified 45,207, 35,894, and 64,679 ATAC-seq peaks in these three conditions, respectively. Quality control analyses indicated high data quality (Supplemental Table S3), strong agreement between experimental replicates (Supplemental Fig. corresponded strongly with previously published datasets performed in relevant cell types, including DNase-seq and ChIP-seq peaks for H3K27ac histone marks (Supplemental Table S7). We examined the impact of EBNA2 on chromatin accessibility by performing differential analysis using MAnorm (Shao et al. 2012) and the IDR approach (Landt et al. 2012) to discover highly reproducible changes in ATAC-seq peaks (see Methods), identifying 1,547 and 690 EBNA2-dependent open and closed chromatin regions, respectively (Fig. 3, Supplemental Table S8). As expected, EBNA2 ChIP-seq peaks in Ramos EBV^EBNA2+^ cells were highly enriched within both EBNA2-dependent open chromatin accessibility gains (85 peaks; 12.4-fold enrichment; adjusted *p*-value: 4.06E^−104^) and EBNA2-dependent chromatin accessibility losses (321 peaks; 96.0-fold enrichment; adjusted *p*-value: 5.48xE^−212^) (Supplemental Table S6). For example, we identified strong EBNA2 ChIP-seq peaks and highly EBNA2-dependent open chromatin in the promoter region of *NAALADL2-AS2*, an uncharacterized antisense long non-coding RNA that is the most highly up-regulated EBNA2 DEG (Fig. 4A).

**Figure 3:**
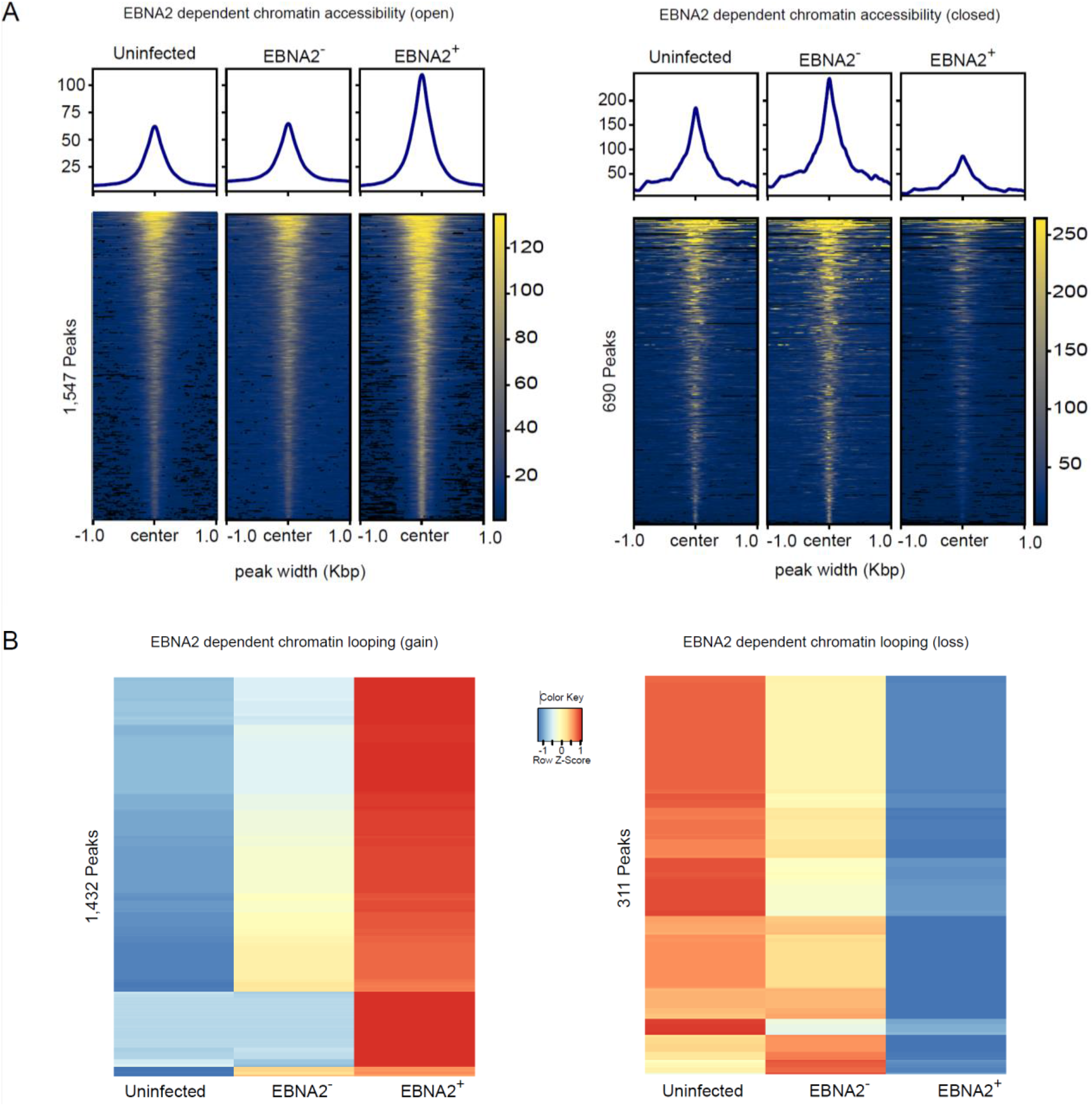
EBNA2 dependent chromatin accessibility and chromatin looping. (A) EBNA2-dependent open chromatin regions and EBNA2-dependent closed chromatin regions are depicted on the left and right, respectively. Values in the heatmaps indicate normalized read counts per genomic region and were generated using the computeMatrix tool in the deepTools package (Ramirez et al. 2016). (B) EBNA2-dependent chromatin loop gains (left) and losses (right). Values in the heatmaps indicate the normalized relative Z-score of the FPKMs across each row (i.e., the default normalization method in the R ‘heatmap’ function).

**Figure 4.**
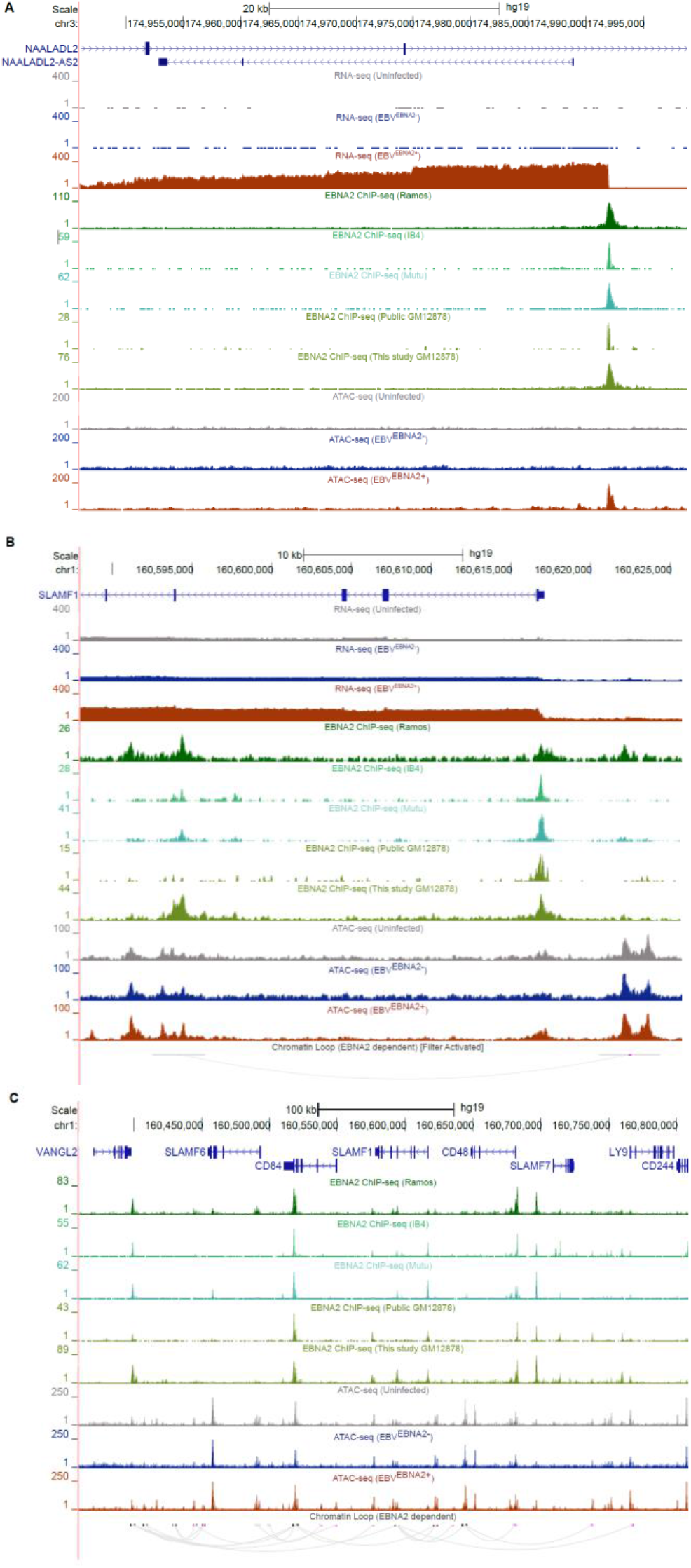
EBNA2-dependent alteration of the human chromatin landscape. (*A*) EBNA2 binding and EBNA2-dependent chromatin opening at the promoter of the upregulated EBNA2 DEG *NAALADL2-AS2*, the most upregulated EBNA2 DEG (UCSC Genome Browser screenshot (hg19)). *(B,C)* EBNA2-dependent alteration of chromatin looping at the *SLAMF1* locus. *(B)* EBNA2-dependent looping to the promoter of the EBNA2 DEG *SLAMF1.* Loops outside of the window are not shown. (*C*) Extensive EBNA2-dependent rewiring of the chromatin looping landscape at the *SLAMF1* locus.

We next tested the hypothesis that EBNA2 up-regulates genes by opening chromatin and down-regulates genes by closing chromatin. We used RELI to examine the significance of the intersection of EBNA2-dependent ATAC-seq peaks, and genomic loci harboring EBNA2 DEGs. As expected, these analyses revealed that EBNA2-dependent open chromatin regions tend to fall proximal (within 5kb of the TSS) to upregulated EBNA2 DEGs (17.4-fold enriched, adjusted *p*-value: 4.30E^−68^), but not proximal to downregulated EBNA2 DEGs (adjusted *p*-value = 1). Likewise, EBNA2-dependent closed chromatin showed the opposite effect, with enrichment for downregulated EBNA2 DEGs (21.5-fold enriched, adjusted *p*-value: 2.77E^−50^) but not upregulated EBNA2 DEGs (adjusted *p*-value = 1) (Supplemental Table S6). Similar results were obtained for distal (100k window) regions of EBNA2 DEGs. Intriguingly, particular TF binding motifs are preferentially enriched within EBNA2-dependent open vs. closed chromatin regions. For example, Ets-like (including PU.1), TCF, and E-box motifs are much more strongly enriched in open regions, whereas ZBED, SOX, and several C2H2 zinc finger motifs are much more strongly enriched in closed regions (Supplemental Table S5). These results collectively reveal an important role for EBNA2 in genome-wide alteration of human chromatin accessibility.

### EBNA2 extensively alters the human chromatin looping landscape

Previous reports support a role for EBNA2 in regulating the three-dimensional structure of chromatin looping (Zhao et al. 2011; McClellan et al. 2013; Jiang et al. 2017). Yet, EBNA2’s roles in human chromatin looping have not been examined genome-wide. Our integrative analysis of EBNA2 DEGs, EBNA2 ChIP-seq peaks, and EBNA2-specific ATAC-seq peaks revealed significant enrichment at these loci for chromatin looping factors such as CTCF, RAD21, and YY1 (Supplemental Table S4), further supporting a possible role for EBNA2 in chromatin looping alteration. To further elucidate the impact of EBNA2 on chromatin looping across the human genome, we next performed HiChIP-seq with an antibody against H3K27ac in the uninfected, EBV^EBNA2−^, and EBV^EBNA2+^ conditions (see Methods).

Analysis of the HiChIP-seq data revealed 93,354, 131,296, and 136,689 chromatin looping interactions in the three conditions, respectively. QC analyses using HiC-Pro (Servant et al. 2015) indicate that our HiChIP-seq experiments are of high quality. The final set of unique valid interaction pairs were between 41.4% and 51.9% of the total sequenced pairs and the number of trans interactions were between 8.8% and 9.9% of the sequenced pairs (Supplemental Table S3), similar to the results obtained in the original HiChIP study (Mumbach et al. 2016). Likewise, the experimental replicates are in high agreement with one another (Supplemental Fig. S4). The full quantification of each chromatin looping event and comparison of these events between conditions are provided in Supplemental Table S9. As expected, EBV^EBNA2+^ looping interactions significantly align (3.8-fold enrichment, *p*-value: 4.08E^−93^, see Methods) with data from a previously published Hi-C experiment performed in GM12878 cells (Rao et al. 2014). Furthermore, most of the EBV^EBNA2+^ HiChIP-seq peaks coincide with publicly available H3K27ac marks, as expected (for example, 78% of HiChIP-seq peaks have these marks in GM19203 cells, 4.0-fold enrichment, adjusted p-value < 1E^−300^) (Supplemental Table S10). Also as expected, ChIP-seq peaks obtained from relevant cell types for POL2RA and chromatin looping factors such as CTCF, RAD21, and YY1 significantly intersected with HiChIP-seq loop anchors, along with active chromatin marks and DNase peaks (Supplemental Table S10). Collectively, these results indicate that our HiChIP-seq data are of high quality.

We next used the HiChIP-seq data to identify EBNA2-dependent differential chromatin looping events (see Methods), identifying 1,432 and 311 looping events that are significantly stronger or weaker in the presence of EBNA2, respectively (Fig. 3B, Supplemental Table S9). As expected, EBNA2-dependent “loop gains” intersect much more significantly with the promoters of upregulated EBNA2 DEGs than with the promoters of downregulated EBNA2 DEGs. Likewise, EBNA2-dependent “loop losses” significantly intersect with downregulated EBNA2 DEGs, but not upregulated DEGs (Supplemental Table S11). These findings support a model where EBNA2-induced promoter interactions increase gene expression levels, while EBNA2-induced loss of promoter interactions decreases gene expression levels. In total, 102 newly formed EBNA2-dependent loops fall within the promoters of 70 EBNA2 DEGs (45 upregulated and 25 downregulated genes) (Supplemental Table S11). EBNA2-dependent gains and losses in chromatin looping were both highly enriched for EBNA2 ChIP-seq peaks (12.0-fold enrichment; adjusted *p*-value: 4.77E^−175^, and 10.9-fold enrichment; adjusted *p*-value: 3.04E^−102^, EBNA2-dependent chromatin loop gains and losses respectively) (Supplemental Table S11). EBNA2-dependent gains in chromatin looping were enriched for EBNA2-dependent chromatin accessibility gains (4.8-fold enrichment; adjusted *p*-value: 1.92E^−26^) but not losses (*p*-value > 0.05). Likewise, EBNA2-dependent losses in chromatin looping were enriched for chromatin accessibility losses (8.8-fold enrichment; adjusted *p*-value: 5.01E^−13^), but not gains (*p*-value > 0.05) (Supplemental Table S11). 58 EBNA2-dependent ATAC-seq peaks are in the promoters of EBNA2 DEGs (TSS +/- 5kb). 43 EBNA2-dependent ATAC-seq peaks are within Ramos

EBV^EBNA2+^ loop anchors that loop to within 20,000 bp of an EBNA2 DEG. Collectively, these results indicate strong agreement between EBNA2-dependent gene expression, binding, chromatin accessibility, and chromatin looping events. For example, EBNA2 binds within the *SLAMF1* gene body, resulting in an EBNA2-dependent loop to the promoter of *SLAMF1*, which is one of the most strongly up-regulated EBNA2-dependent genes (Fig. 4B). Remarkably, the *SLAMF1* locus contains dozens of robust EBNA2-dependent chromatin looping and accessibility events involving multiple genes (Fig. 4C), revealing extensive, EBNA2-dependent rewiring of the chromatin threedimensional landscape at this locus.

### EBNA2-dependent mechanisms significantly coincide with autoimmune disease risk loci

Previous studies (Ricigliano et al. 2015; Harley et al. 2018) have nominated an important role for EBNA2 in autoimmune and other human diseases. We therefore systematically compared the genomic locations of EBNA2-dependent mechanisms to the locations of genetic risk variants obtained from 172 published GWAS datasets (see Methods). Strikingly, the genomic regions surrounding EBNA2 DEGs were enriched for many of the same autoimmune diseases we previously identified based on intersection of GWAS signal with EBNA2 ChIP-seq peaks (Harley et al. 2018). Specifically, 65 of the 421 EBNA2 DEGs (39 up-regulated, 26 down-regulated) have GWAS signal for autoimmune disorders within 100 kilobases of their transcription start site (Fig. 5A, Supplemental Table S12). Among these, 20 and 19 EBNA2 DEGs are located within SLE and MS-associated loci, respectively. In contrast, EBNA2-independent EBV DEGs (i.e, genes with higher or lower expression in both EBV^EBNA2+^ and EBV^EBNA2−^ conditions compared to uninfected) do not coincide with autoimmune disease risk loci (Supplemental Table S12), suggesting an EBNA2-specific (as opposed to EBV-specific) role in autoimmune gene regulatory mechanisms.

**Figure 5.**
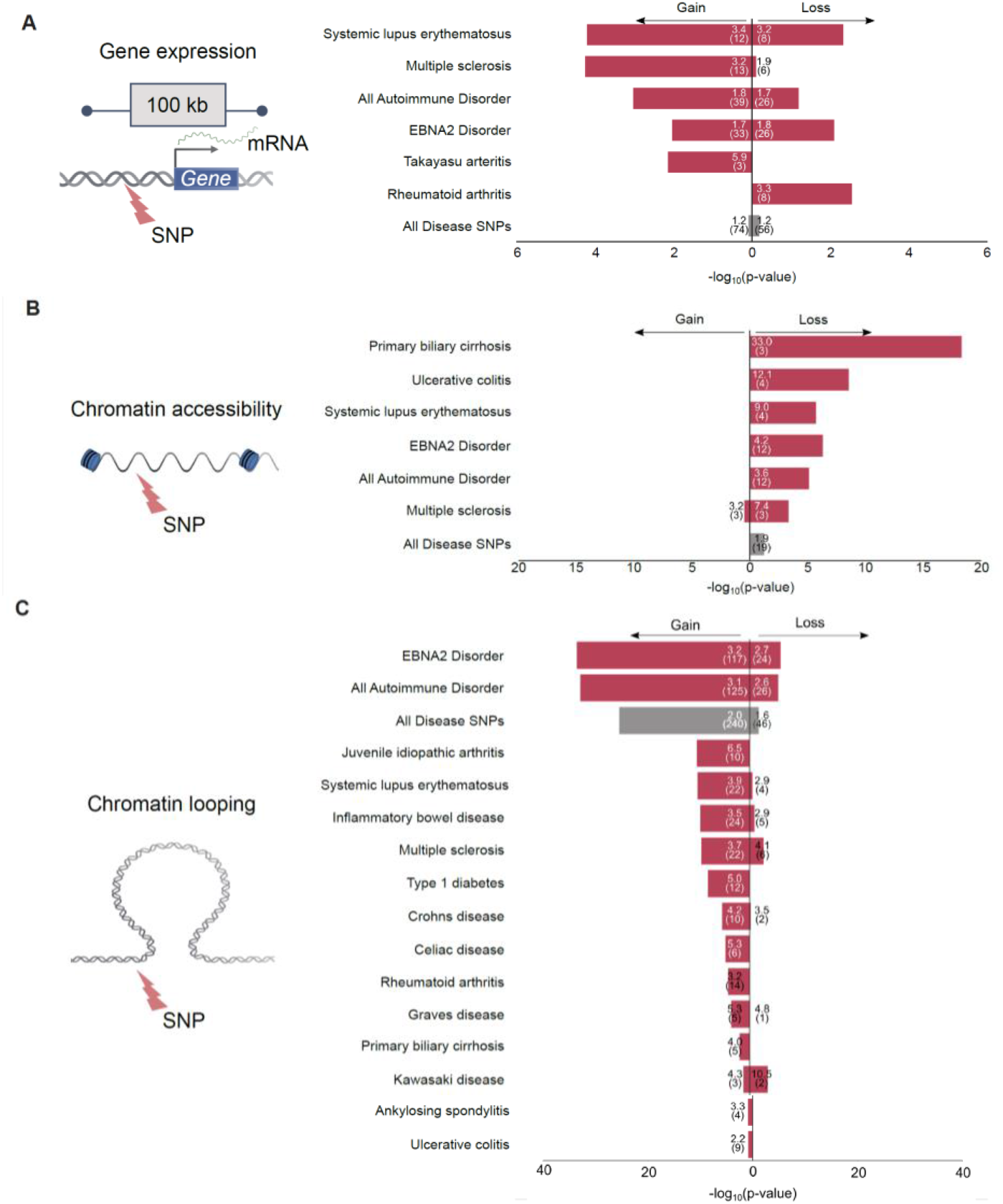
Intersection of EBNA2-dependent gene regulatory mechanisms and disease-associated genetic variants. . The bar plots indicate the significance of the intersection (RELI negative log10 adjusted *p*-value). For all analyses, 19 autoimmune diseases were individually tested, along with a set of variants from all 19 autoimmune diseases (“All Autoimmune Disorders”), a set containing variants from the nine “EBNA2 Disorders” from our previous study (Harley et al. 2018), and a set containing 176 diseases and phenotypes (“All Disease SNPs”). Only diseases with at least one significant result (3 or more overlaps, corrected *p*-value < 0.05) are shown. Autoimmune diseases are indicated with red bars. Fold-enrichment and number of overlaps are indicated inside the bars. (*A*) Significant intersection between EBNA2-dependent differentially expressed genes and disease-associated variants (‘Gain’: upregulated genes; ‘Loss’: downregulated genes). (B) Significant intersection between EBNA2-dependent chromatin accessibility and disease-associated variants (‘Gain’: newly opened chromatin; ‘Loss’: newly closed chromatin, relative to uninfected). (C) Significant intersection between EBNA2-dependent chromatin looping and disease-associated variants (‘Gain’: new looping events; ‘Loss’: loss of looping events, relative to uninfected).

Next, we inspected EBNA2-dependent chromatin accessibility regions for disease enrichment. Surprisingly, despite the vast changes in chromatin accessibility orchestrated by EBNA2, we did not identify significant intersection between EBNA2-dependent chromatin opening events and autoimmune-associated variants (Fig. 5B, Supplemental Table S12). Rather, we observed enrichment for autoimmune variants within EBNA2-dependent chromatin closing events (3.6-fold enrichment, adjusted *p*-value: 7.73E^−6^) (Fig. 5B, Supplemental Table S12). These results prompted us to further examine the relationship between chromatin accessibility and autoimmune risk loci. To this end, we examined the significance of the intersection between autoimmune associated variants and several types of ATAC-seq peaks: 1) constitutively open chromatin regions (i.e., peaks shared in uninfected, EBV^EBNA2+^, and EBV^EBNA2−^ conditions); 2) EBV-dependent peaks (gains and losses); 3) EBV^EBNA2−^-dependent peaks (gains and losses); and 4) EBV^EBNA2+^-dependent peaks (gains and losses). These analyses revealed significant intersection between autoimmune variants and regions of the genome that are constitutively open in B cells (Supplemental Table S13). These results indicate that autoimmune risk variants also concentrate in regions of the genome that are already accessible for the binding of EBNA2 and other proteins prior to infection. Collectively, these analyses reveal that EBNA2-dependent opening of chromatin is not a significant component of autoimmune risk loci; however, EBNA2-dependent chromatin closing, and accessible chromatin prior to EBV infection, are.

Finally, we investigated the relationship between EBNA2-altered chromatin looping interactions and autoimmune disease risk loci. This analysis revealed highly significant intersection between EBNA2-induced changes to chromatin looping and autoimmune disease risk loci. In particular, 125 newly established EBNA2-dependent chromatin loop anchors intersect autoimmune-associated variants (3.1-fold enrichment; adjusted *p*-value: 9.07E^−33^) (Fig. 5C, Supplemental Table S12). EBNA2-dependent loss of chromatin looping also shows significant intersection with autoimmune variants, albeit to a much lesser degree (Fig. 5C, Supplemental Table S12). Collectively, these data indicate that EBNA2-dependent alteration of long-range chromatin interactions is highly associated with autoimmune disease-associated genetic variants, revealing a key role for EBNA2-altered chromatin interactions in autoimmune disease etiology.

### Allele-dependent EBNA2 mechanisms at autoimmune-disease risk loci

The previous analyses revealed that EBNA2-dependent chromatin alterations are significantly enriched for autoimmune-associated genetic variants. We therefore next used our MARIO pipeline (Harley et al. 2018) to systematically identify autoimmune risk allele-dependent EBNA2 binding events that coincide with these EBNA2-dependent mechanisms (see Methods). In brief, MARIO identifies genotype-dependent (allelic) functional genomic data (i.e. read imbalance) at genomic locations where the assayed cell contains both the reference and non-reference alleles (i.e., the genotype of the cell must be heterozygous for that specific polymorphism).

Using this approach, we discovered 32 instances of allele-dependent EBNA2 binding at autoimmune risk variants (8 in EBV^EBNA2+^ Ramos, and 24 in GM12878 cells) (Supplemental Table S14), including validation in Ramos cells of the allele-dependent EBNA2 binding we previously observed for rs3794102 at the *CD44* locus in Mutu cells (Harley et al. 2018). For example, we identified strong EBNA2 allele-dependent binding in EBV^EBNA2+^ Ramos cells to an MS-associated variant (rs1250567) located in the *ZMIZ1* locus (Fig. 6). This region loops to the promoter of the short isoform of *ZMIZ1* in EBV^EBNA2+^ Ramos cells. rs1250567 is a strong eQTL for *ZMIZ1* in both EBV-immortalized lymphoblast cell lines (p= 1.76E^−4^) and whole blood (p= 2.21E^−4^) (eQTL Catalogue: https://www.biorxiv.org/content/10.1101/2020.01.29.924266v1) (Supplemental Table S15). *ZMIZ1* expression levels are three-fold lower in EBV^EBNA2+^ Ramos cells compared to uninfected Ramos cells, consistent with a previous report by Fewing et al. describing decreased ZMIZ1 protein expression in MS patient blood samples (Fewings et al. 2017). Collectively, our results at the *ZMIZ1* locus reveal EBNA2 and autoimmune risk allele-dependent mechanisms possibly underlying the established roles played by genetics and the environment in autoimmune diseases.

**Figure 6.**
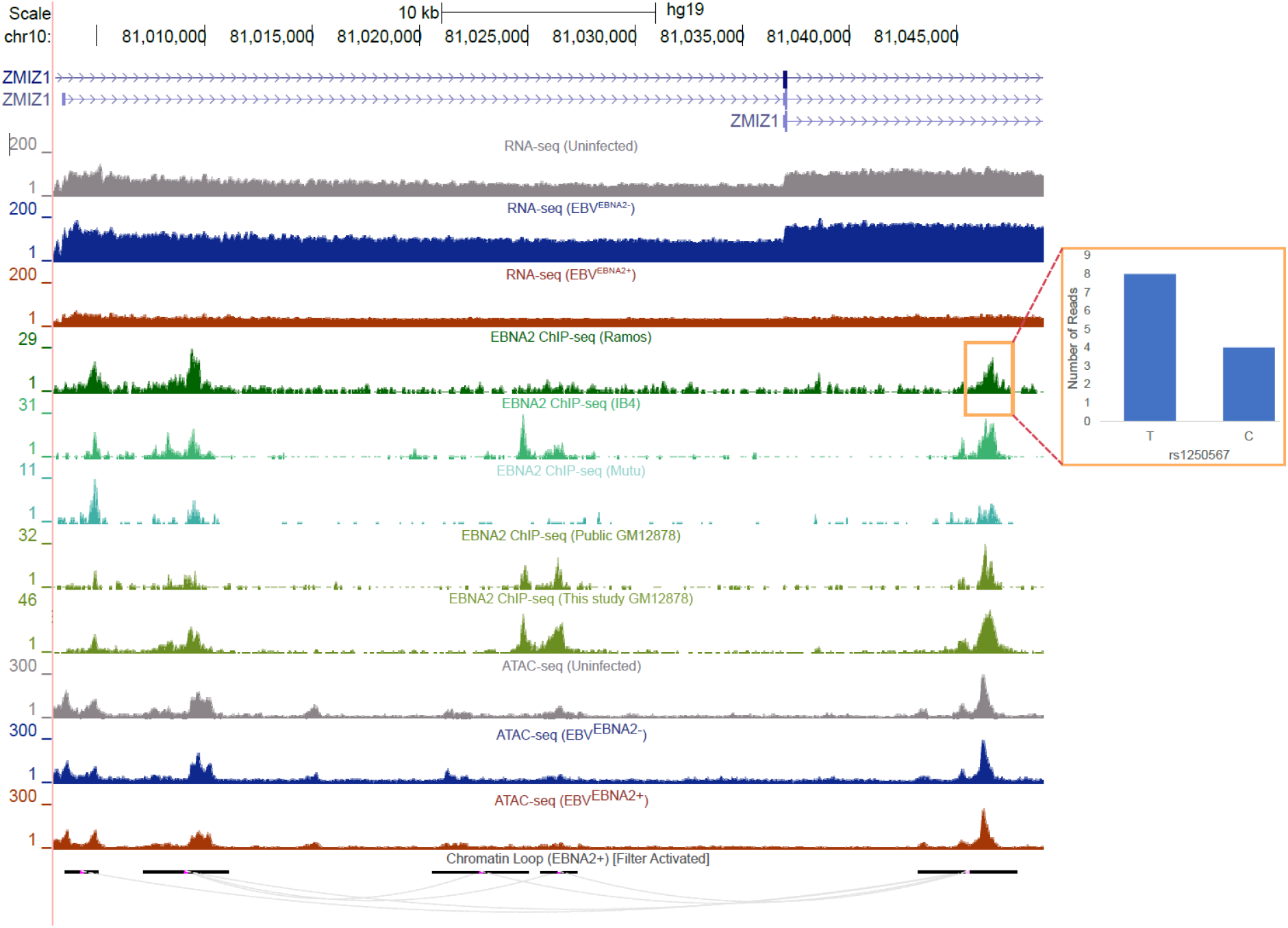
Allele-dependent and EBNA2-dependent mechanisms at an autoimmune-associated genetic variant. . UCSC Genome Browser shots (hg19) depicting EBNA2-based mechanisms at the *ZMIZ1* locus. Loops outside of the window are not shown (filtered). Red box indicates the region where EBNA2 binds in an allele-dependent manner. The ratio of reads between alleles is shown as a bar plot. See text for details.

In total, 633 unique autoimmune-associated genetic variants can be implicated in at least one EBNA2-dependent mechanism (Supplemental Table S16). Among these, 24 are involved in allele-dependent EBNA2 binding, 41 are located within the promoters (TSS +/- 500bp) of EBNA2 DEGs, 46 are located within EBNA2-dependent chromatin accessibility regions, and 539 are located within EBNA2-dependent chromatin looping events. Collectively, these data implicate EBNA2 in multiple types of autoimmune disease gene regulatory mechanisms.

## Discussion

In this study, we examined the mechanisms by which an EBV-encoded transcriptional regulator, EBNA2, modulates human gene regulatory programs through genome-wide perturbation of the human chromatin landscape. Our findings show that EBNA2 affects cellular gene regulation using three of the same mechanisms a host regulatory protein might use: 1) interacting with promoters and enhancers; 2) altering chromatin accessibility; and 3) forming new chromatin 3D interactions. We demonstrate that many of these mechanisms significantly intersect autoimmune-associated genetic variants, and identify multiple EBNA2 interactions that are autoimmune risk alleledependent.

Although it was not the focus of this study, we also identify many “EBNA2-independent” genes and regulatory elements in the human genome, which are stronger/weaker in both EBV^EBNA2−^ and EBV^EBNA2+^ cells compared to uninfected cells. These genes and elements are altered by EBV infection independently of EBNA2, and hence might involve other EBV-based regulatory molecules, such as Zta, Rta, or the EBNA3 proteins (Liu et al. 2020).

It has been extensively reported that multiple human diseases have a strong EBV-based component (Bray et al. 1983; Serafini et al. 2007; Pender and Burrows 2014) or EBNA2 component (Ricigliano et al. 2015; Harley et al. 2018). In this study, by controlling both EBV infection and the presence of EBNA2 in the EBV-infected cells, we not only corroborate these previous findings, but also identify additional gene targets and mechanisms by which EBV and EBNA2 might affect the course of these diseases. In particular, we observe significant intersection between autoimmune-associated genetic variants and EBNA2-dependent modulation of human gene expression, chromatin accessibility, and chromatin looping interactions (Fig. 5).

Previous studies have demonstrated a key role for EBNA2 in multiple biological processes, including B cell transformation (Saha and Robertson 2019), interferon regulation (Kanda et al. 1992), and NF-kB regulation (Kim et al. 2017). In this study, we find that several genes involved in leukocyte cell-cell adhesion display EBNA2-dependent gene expression levels and regulatory mechanisms, including *NAALADL2-AS2, CD80, SLAMF1*, and *ZMIZ1.* Importantly, several of these interactions are autoimmune disease risk-allele dependent. EBV infection is known to alter the expression of adhesion molecules and receptors (Zhao et al. 2006; Shannon-Lowe and Rowe 2011; Grossman et al. 2017). In particular, Fewing et al. reported a negative correlation between

*ZMIZ1* expression and EBV antigen levels (Fewings et al. 2017), consistent with the EBNA2-dependent decrease in *ZMIZ1* expression that we observe. Further, previous studies have reported elevated expression levels of adhesion molecules in SLE patients (Funauchi et al. 1993; Egerer et al. 2000) and MS patients (Elovaara et al. 2000). Together, these results support a model where EBNA2 affects autoimmune disease onset and/or progression through the alteration of cell-cell adhesion-related gene regulatory programs.

In addition to possible EBNA2 roles in the regulation of cell-cell adhesion genes, we identified numerous strongly EBNA2-dependent genes. Consistent with the results from Wang et al., *NAALADL2-AS2* was the gene with the highest degree of EBNA2-dependent upregulation. *NAALADL2-AS2* is a long non-coding RNA expressed on the opposite strand of *NAALADL2*, with minimal expression across the immune compartment with the exception of naïve B cells (Shay and Kang 2013). Another strongly upregulated gene we detected is Troponin T3, Fast Skeletal Type *(TNNT3)*, which is expressed across lymphocyte subsets and has been shown to interact with the EBNA2 co-activator EBNA-LP (Kempkes and Ling 2015; Rouillard et al. 2016). Solute Carrier Family 25 Member 24 (SLC25A24) is a calcium binding carrier protein with strong EBNA2-dependent decreased gene expression. A previous study of patients with the EBV-associated Burkett’s lymphoma found that patient death was associated with SLC25A24 downregulation when the tumor was EBV-positive (Kaymaz et al. 2017).

We observed widespread EBNA2-dependent chromatin looping events at a genomic locus encoding multiple members of the signaling lymphocyte activation molecule (SLAM) family, including *CD84, SLAMF1*, and *LY9.* SLAM family genes were previously reported to have EBV-dependent gene expression patterns (Wang et al. 2019), with *SLAMF1* in particular being among the most strongly up-regulated EBNA2 target genes (Maier et al. 2006). SLAM family genes play key roles in several immunological processes, including humoral responses (Ma et al. 2007), development and maintenance of immune system function (Schwartzberg et al. 2009), and cell adhesion (reviewed in (Cannons et al. 2011)). Our new results establish a possible role for EBNA2 in the regulation of the SLAM-mediated autoimmune-related immune response.

At the *ZMIZ1* locus, we present genotype-dependent binding of EBNA2 at a disease risk variant, looping of the genetic variant to the “short isoform” promoter of *ZMIZ1*, EBNA2-dependent expression of the *ZMIZ1* “short isoform”, and allele-dependent expression of *ZMIZ1* as a function of the disease risk variant genotype. While these results are consistent with our conclusion of EBNA2-dependent allelic expression of *ZMIZ1*, it is possible that additional gene regulatory mechanisms might also mediate the expression of *ZMIZ1* and contribute to disease.

A limitation of our study design is that P3HR-1 and B95.8 strains are different genetic isolates of EBV. Therefore, there are further genetic differences between the strains in addition to the EBNA2 deletion. For example, the last two exons (45 amino acids) of EBNA-LP are also deleted in the P3HR-1 virus. However, EBV mutants with these two exons deleted can still transform primary B cells, albeit at lower frequencies (Mannick et al. 1991). Further, P3HR-1 is a well-established model system that has been widely used as an EBNA2-null strain in the field of virology for decades (Murray et al. 1988; Zimber-Strobl et al. 1991; Lee et al. 2002; Jiang et al. 2017).

In summary, our findings reveal an important role for the EBV-encoded EBNA2 regulatory protein in multiple mechanisms that ultimately affect human gene expression levels. Several of these mechanisms are allele-dependent at variants associated with autoimmune diseases such as MS and SLE. It is possible that other viral transcriptional regulators play similar mechanistic roles in other diseases. Future studies will deepen our knowledge of the mechanisms underlying virus-host interactions, and ultimately provide both a rationale and a foundation for therapeutic approaches targeting these interactions.

## Methods

### EBV infection of Ramos B cells

Wild-type B95.8 EBV (EBV^EBNA2+^) and P3HR-1 EBV lacking EBNA2 (EBV^EBNA2−^) were prepared from B95.8 and P3HR-1 cell supernatants respectively, and cultured in 10% FBS supplemented RPMI medium 1640 for 2 weeks. Viral suspension was filtered via 0.45 μm Millipore filters and cells were pelleted. The concentrated viral stocks were stored at −80 °C. Ramos cells (EBV Negative, ATCC CRL-1596) were infected ~2 × 10^6^/mL with viral stock based on infection optimization assays and incubated for 4 hours for virus adsorption. After infection, cells were washed and cultured. After 10 passages, we confirmed the infection by morphological changes and the expression of EBNA2 viral protein levels as previously published (Harley et al. 2018).

### RNA-seq

Total RNA was extracted using the mirVANA Isolation Kit (Ambion) from Ramos B cells in each of the three conditions: uninfected Ramos (uninfected), EBNA2-positive EBV (B95.8)-infected Ramos (EBV^EBNA2+^), and EBNA2-negative EBV (P3HR-1)-infected Ramos (EBV^EBNA2−^). RNA was extracted at the same time point for every condition. RNA was depleted of ribosomal RNA using the Ribo-Zero rRNA removal kit and sequenced at Cincinnati Children’s Hospital Medical Center (CCHMC) DNA Sequencing and Genotyping Core Facility. Using the tool FastQC (version: 0.11.2) (http://www.bioinformatics.babraham.ac.uk/projects/fastqc), all data were confirmed to pass all quality control checks, except for adapter sequence contents, which were removed using cutadapt (trimgalore version: 0.4.2)(https://journal.embnet.org/index.php/embnetjournal/article/view/200). RNA-seq reads were aligned to the hg19 (GrCh37) genome build (NCBI) using Spliced Transcripts Alignment to a Reference (STAR, version: 020201) (https://doi.org/10.1093/bioinformatics/bts635). The featureCounts function (Rsubread version: 3.5.3) was used to count reads (Liao et al. 2014) and DESeq2 (version 3.5.3) was used to perform differential gene expression (Love et al. 2014). DESeq2 results were filtered to only include genes with fragments per kilobase per million (FPKM) values greater than one in at least one condition. Differential expression was calculated with thresholds of a greater than 1.5-fold change and an adjusted *p*-value threshold of less than 0.05 (*p*-values were adjusted by Benjamini and Hochberg false discovery rate (FDR)). Gene set enrichment analysis was performed using ToppFun (Chen et al. 2009).

### ChIP-seq

ChIP-seq for EBNA2 was performed in duplicate in Ramos (EBV^EBNA2+^) and GM12878 cell lines. Ramos and GM12878 cells were crosslinked and nuclei were sonicated as described previously (Lu et al. 2015). Cells were incubated in crosslinking solution (1% formaldehyde, 5 mM HEPES [pH 8.0], 10 mM sodium chloride, 0.1 mM EDTA, and 0.05 mM EGTA in RPMI medium 1640 with 10% FBS) and placed on a tube rotator at room temperature for 10 min. To stop the crosslinking, glycine was added to a final concentration of 0.125 M and tubes were placed back on the rotator at room temperature for 5 min. Cells were washed twice with ice-cold PBS, resuspended in lysis buffer 1 (50 mM HEPES [pH 7.5], 140 mM NaCl, 1 mM EDTA, 10% glycerol, 0.25% Triton X-100, and 0.5% NP-40), and incubated for 10 min on ice. Nuclei were harvested after centrifugation at 5,000 rpm for 10 min, resuspended in lysis buffer 2 (10 mM Tris-HCl [pH 8.0], 1 mM EDTA, 200 mM NaCl, and 0.5 mM EGTA), and incubated at room temperature for 10 min. Protease and phosphatase inhibitors were added to both lysis buffers. Nuclei were then resuspended in sonication buffer (10 mM Tris [pH 8.0], 1 mM EDTA, and 0.1% SDS). A S220 focused ultrasonicator (COVARIS) was used to shear chromatin (150- to 500-bp fragments) with 10% duty cycle, 175 peak power, and 200 bursts per cycle for 7 min. A portion of the sonicated chromatin was run on an agarose gel to verify fragment sizes. Sheared chromatin was precleared with 20 μL Dynabeads Protein G (Life Technologies) at 4 °C for 40 min.

Immunoprecipitation of EBNA2-chromatin complexes was performed with an SX-8X IP-STAR compact automated system (Diagenode). Beads conjugated to antibodies against EBNA2 (Abcam; ab90543) were incubated with precleared chromatin at 4°C for 8 hours. The beads were then washed sequentially with wash buffer 1 (10 mM Tris-HCl [pH 7.5], 150 mM NaCl, 1 mM EDTA, 0.1% SDS, 0.1% NaDOC, and 1% Triton X-100), wash buffer 2 (10 mM Tris-HCl [pH 7.6], 400 mM NaCl, 1 mM EDTA, 0.1% SDS, 0.1% NaDOC, and 1% Triton X-100), wash buffer 3 (10 mM Tris-HCl [pH 8.0], 250 mM LiCl, 1 mM EDTA, 0.5% NaDOC, and 0.5% NP-40), and wash buffer 4 (10 mM Tris-HCl [pH 8.0], 1 mM EDTA, and 0.2% Triton X-100). Finally, the beads were resuspended in 10 mM Tris-HCl (pH 7.5) and used to prepare libraries via ChIPmentation (Schmidl et al. 2015).

The resulting libraries were sequenced targeting 100,000,000 unique single-end reads. We performed quality control of raw sequencing reads using FastQC (version: 0.11.2) (http://www.bioinformatics.babraham.ac.uk/projects/fastqc). All data were confirmed to pass all quality control checks (Supplemental Table S3), except for adapter sequence contents, which were removed using cutadapt (trimgalore version: 0.4.2). Alignment of reads to the human genome (build hg19) was performed using Bowtie2 (Langmead and Salzberg 2012). Peaks were called using Model-based Analysis of ChIP-Seq version 2.1.1 (MACS2) (Feng et al. 2012) with the following arguments: -g hs -q 0.01 --broad. These settings were chosen from several tested parameter combinations because they yielded the best TF binding site motif enrichment *p*-values (using HOMER) for EBNA2 binding partner RBPJ. We observed strong agreement between experimental replicates (Supplemental Fig. S4). We thus pooled the reads between the replicates and re-called peaks using MACS2, in order to capture as many EBNA2 binding events as possible. We also obtained publicly available EBNA2 ChIP-seq datasets (GM12878 cells: SRR3101734; IB4 cells: SRR332246; Mutu cells: SRR8783824), and called peaks using MACS2 with the following arguments: -g hs -q 0.01.

### ATAC-seq

ATAC-seq was performed in duplicate in the three Ramos cell conditions. Briefly, transposase Tn5 with adapter sequences was used to cut accessible DNA. These accessible DNA with adaptor sequences were isolated, then libraries were prepared for uninfected, EBV^EBNA2−^, EBV^EBNA+^ Ramos using a standard protocol (Buenrostro et al. 2015). Paired-end sequencing was performed at the CCHMC DNA Sequencing and Genotyping Core Facility. FastQC was used to perform quality control (http://www.bioinformatics.babraham.ac.uk/projects/fastqc), as described above.

ATAC-seq reads were aligned to the human genome (hg19) using Bowtie2 (Langmead and Salzberg 2012) and peaks were called using MACS2 (same version and parameter settings as for our Ramos ChIP-seq) (Feng et al. 2012). We observed strong agreement between experimental replicates (Supplemental Fig. S4). We thus pooled the reads between the replicates and re-called peaks using MACS2, in order to capture as many open chromatin regions as possible. Differential chromatin accessibility was calculated using the MAnorm program (Shao et al. 2012) and bedtools (Quinlan and Hall 2010). First, we determined EBV-dependent open chromatin by calculating EBV^EBNA2+^-unique peaks compared to uninfected peaks (>1.5 fold, *p*-value < 0.05) using MAnorm. Next, to obtain EBNA2-dependent regions, we subtracted EBV^EBNA2−^-unique peaks compared to uninfected using the bedtools subtract command. To identify EBNA2-dependent closed chromatin, we first determined EBV-dependent closed chromatin, by identifying uninfected-unique peaks compared to EBV^EBNA2+^ peaks, then we subtracted the uninfected-unique peaks compared to EBV^EBNA2+^ peaks. Finally, we used the IDR approach (Landt et al. 2012) (IDR threshold of 5%) to identify only highly reproducible changes in ATAC-seq peaks.

### HiChIP-seq

HiChIP-seq libraries were prepared following Mumbach *et al.* (Mumbach et al. 2016). We obtained a single experimental dataset for the uninfected condition, and two experimental replicates for the EBV^EBNA2−^ and EBV^EBNA2+^ conditions. Cells were cross-linked with 1% formaldehyde and lysed with Hi-C lysis buffer. Cross-linked DNA was digested by MboI. DNA ends were filled with biotin-dATP and dCTP, dTTP, dGTP and then ligated by T4 DNA ligase. After ligation, DNA was sonicated using a Covaris M220. Fragmented DNA was then diluted 10 times with ChIP dilution buffer and protein-DNA complexes were captured using an H3K27ac antibody (Abcam AB4729). Next, protein-DNA complexes were further captured by protein A beads and eluted. DNA was purified after reverse crosslinking. Biotin dATP-labeled DNA was captured with Streptavidin C-1 beads and PCR amplified using Phusion HF (New England Biosciences) and 1 μL of each Nextera forward primer (Ad1_noMX) and Nextera reverse primer (Ad2.X). The libraries were sequenced using Illumina Nextseq (2×75bp).

We used HiC-pro (version: 2.11.0) to align reads and identify the Hi-C contact map (Servant et al. 2015). HiC-pro was also used to perform quality control (Supplemental Table S3). We then integrated the aligned Hi-C data with the ChIP-seq data to correct the background, perform restriction site bias modeling and obtain looping information using hichipper (version: 0.7.3) (http://aryeelab.org/hichipper). We observed strong agreement between experimental replicates (Supplemental Fig. S4). We thus pooled the reads between the replicates and re-analyzed the data, as described above. We identified differential looping events using the diffloop Bioconductor R package (1.6.0) (Lareau and Aryee 2018). Briefly, to calculate EBNA2-dependent differential looping, we first determined EBV-dependent chromatin looping by comparing EBV^EBNA2+^ looping events to uninfected looping events using diffloop (>1.5 fold change, *p*-value < 0.05). Likewise, we also compared EBV^EBNA2−^ looping events to uninfected looping events using the same cutoffs. Then to identify EBNA2-dependent looping events, we applied diffloop again, comparing EBV^EBNA2+^-dependent looping events to EBV^EBNA2−^-dependent looping events, using the same cutoffs.

We calculated the significance of the intersection between chromatin looping events in EBV^EBNA2+^ conditions using looping data from GM12878 cells (Rao et al. 2014), using a permutation test. Specifically, we randomized EBV^EBNA2+^ loop coordinates, using 500 iterations of permutation. Permutations were performed by randomly sampling a gene promoter, placing one end of the loop in that promoter, and the other end of the loop the same distance away from that promoter as the “real” loop. This randomization procedure thus controls both for the looping distance and the propensity for loops to have higher densities near gene promoters. We then calculated a z-score by comparing the distribution of randomized intersection counts to actual intersection counts.

### Estimation of the significance of intersected genomic coordinates using RELI

We used the RELI method to calculate the significance of intersection between the genomic coordinates of two or more datasets. In brief, RELI calculates the overlap between the input genomic coordinates and a library of ChIP-seq derived genomic coordinates. It then permutes the input coordinates to calculate the overlap between randomized coordinates and the library coordinates. After 2,000 iterations, RELI then compares the randomized overlap to the actual overlap, and calculates the significance of this overlap (Harley et al. 2018). We used a RELI library containing 1,544 TF datasets, 2,455 Non-TF datasets, and disease-associated genetic variants (through GWAS) from 172 diseases (Harley et al. 2018). Search windows (100 kb for distal, 5kb for promoter) were padded from the transcription start site (TSS) coordinates of EBNA2 DEGs. We ran RELI (null model: OpenChrom) with padded DEGs, EBNA2 ChIP-seq, EBNA2-dependent chromatin, and EBNA2 looping interactions. RELI output was filtered based on the number of overlaps (greater than 3) and significance (adjusted *p*-value < 0.05, Bonferroni correction), following our standard practices.

### Identification of allele-dependent sequencing reads using MARIO

Allele-dependent behavior was identified in sequencing reads using the MARIO pipeline (Harley et al. 2018). Briefly, the MARIO pipeline identifies allele-dependent behavior by weighing 1) the imbalance between the number of reads that are mapped to each allele, 2) the total number of reads mapped at each variant, and 3) the number of and the consistency of available replicates. These variables are combined into a single Allelic Reproducibility Score (ARS), which reflects the degree of allelic behavior observed for the given heterozygous variant in the given dataset. To identify heterozygous variants, we used genotyping array data for Ramos cells (Harley et al. 2018) and GM12878 cells (dbGaP ID: phs001989.v1). Imputation was performed using the impute2 program (Howie et al. 2009). MARIO ARS values exceeding 0.4 were considered to be allelic, following our previous study (Harley et al. 2018).

### TF DNA binding motif enrichment analysis

We used the HOMER software package (Heinz et al. 2010) to calculate TF DNA binding site motif enrichment. We used a modified version of HOMER that incorporates human motifs obtained from CisBP build 2.0 (Weirauch et al. 2014; Lambert et al. 2019).

### GWAS dataset curation

We obtained GWAS data from multiple studies from the NHGRI-EBI GWAS Catalog (version GWAS_catalog_v1.0.2-associations_e96_r2019-05-03) (Buniello et al. 2019). A genome wide significance cutoff of 5 × 10^−8^ was used to establish the statistical significance of a variant and its association to a given disease or trait. After filtering for genome wide association, variants were grouped based on the disease or trait reported in the publication as well as the reported ancestries. For each disease/trait, independent loci were identified using LD-based pruning in Plink (window size 300,000 kb, SNP shift size 100,000 kb, and r2 < 0.2).

## Data and code access

USCC Genome Browser session:

https://genome.ucsc.edu/s/parz1z/EBNA2 Ramos Chromatin paper

GEO accession (RNA-seq, ChIP-seq, ATAC-seq, HiChIP-seq):

GSE148396

dbGAP (genotyping):

phs001989.v1

Published data sets (EBNA2 ChIP-seq):

- GM12878 cells: SRR3101734

- IB4 cells: SRR332246

- Mutu cells: SRR8783824

RELI source code: https://github.com/WeirauchLab/RELI

MARIO source code: https://github.com/WeirauchLab/MARIO

## Acknowledgments

The authors thank Artem Barski for maintaining robot facilities for ChIP-seq experiments. The authors thank Kevin Ernst for critical input on technical issues and computational support including servers, software packages, and data management. The authors also thank Mario Pujato (AstraZeneca), Xiaoting Chen, and Xiaoming Lu for insights related to data analysis. Finally, the authors acknowledge that the cartoons from Figure 1 and Figure 5 were created using BioRender.com.

## Funding sources

This study was supported by NIH grants R01 HG010730, R01 NS099068 and R01 GM055479 (M.T.W.); R01 AR073228 (L.C.K. and M.T.W.); R01 DK107502 and P30 AR070549 (L.C.K.); and U01 AI130830, U01 HG008666, R01 AI024717, R01 AI148276, and I01 BX001834 (J.B.H.). This study was further supported by funding from the Ohio Supercomputing Center (M.T.W.), along with funds from Cincinnati Children’s Research Foundation ARC and CpG awards (M.T.W., L.C.K., and J.B.H.) and a CCRF Endowed Scholar Award (M.T.W.)

## Notes

### Competing Interest Statement

The authors have declared no competing interest.

