## Supplemental Figure S1 for "Epstein-Barr virus nuclear antigen 2 (EBNA2) extensively rewires the human chromatin landscape at autoimmune risk loci"

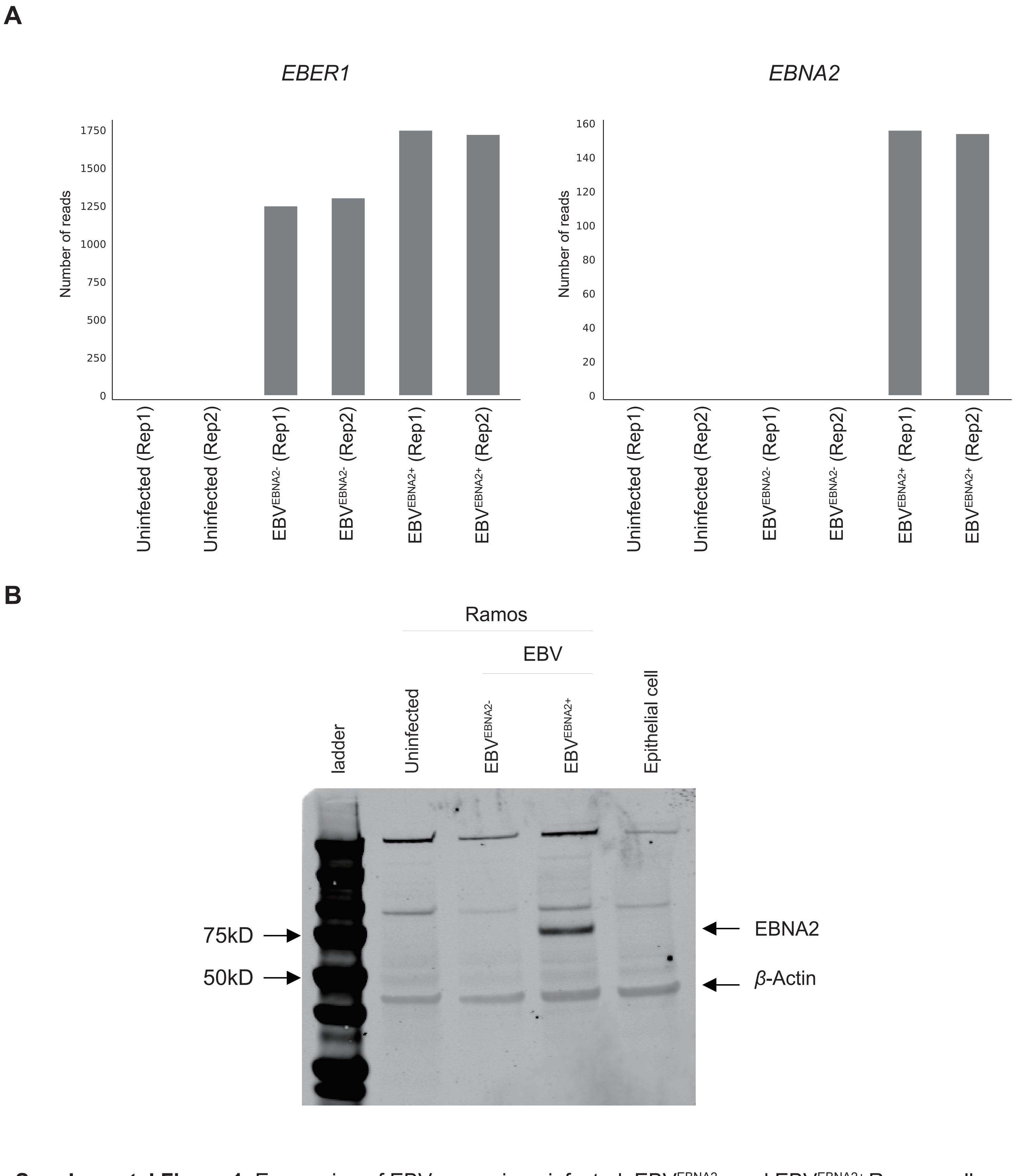


**Supplemental Figure 1**. Expression of EBV genes in uninfected, EBV$^{EBNA2-}$, and EBV$^{EBNA2+}$ Ramos cells. (*A*) Gene expression (read count) of *EBER1* and *EBNA2* among infection types. (*B*) Western blot showing differential EBNA2 expression among infection types.
