## Supplemental Figure S2 for "Epstein-Barr virus nuclear antigen 2 (EBNA2) extensively rewires the human chromatin landscape at autoimmune risk loci"

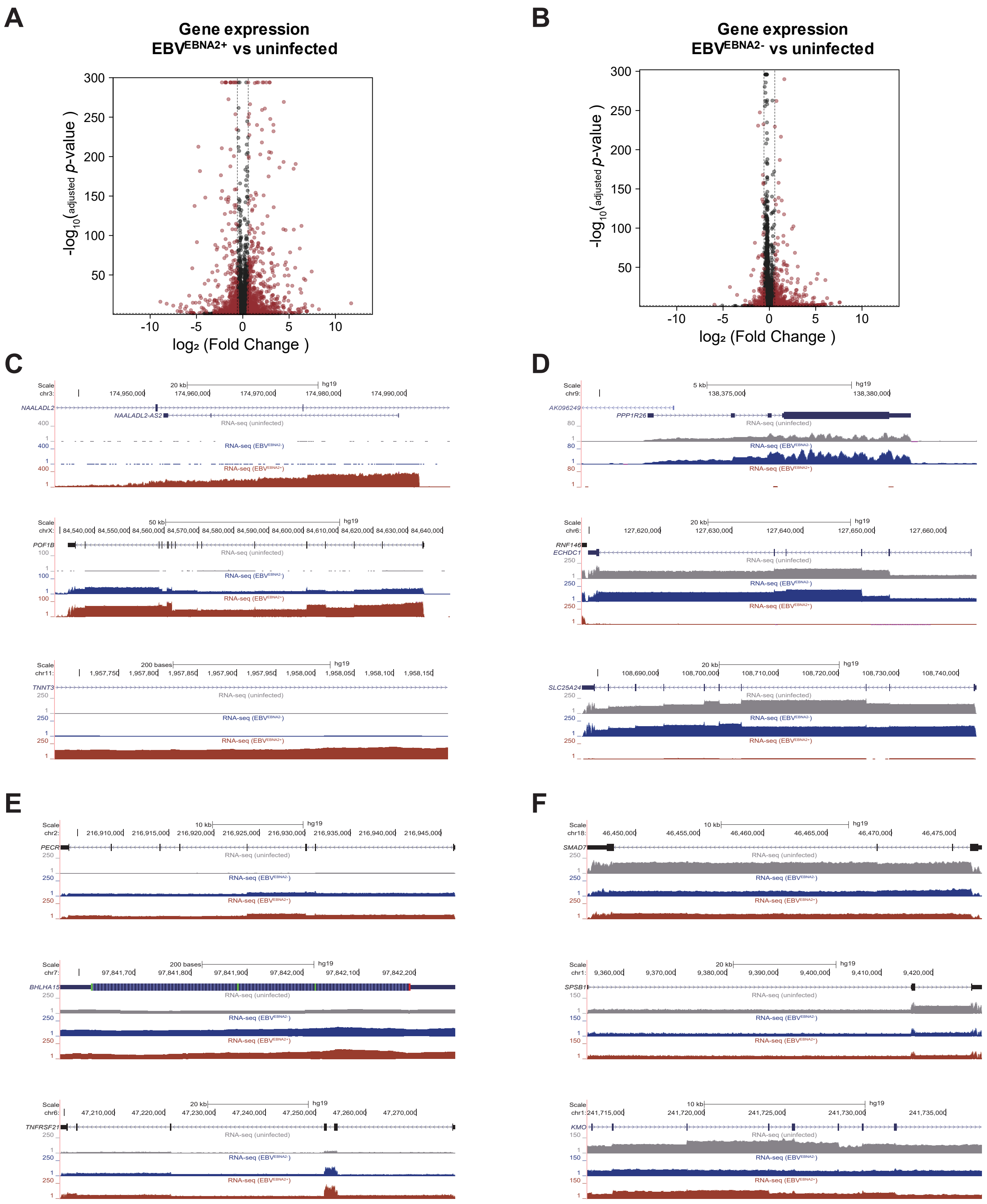

**Supplemental Figure 2**. EBNA2-dependent human gene expression in Ramos B cells. (*A*,*B*). Volcano plots showing differential human gene expression upon infection with EBV$^{EBNA2+}$ (*A*) and EBV$^{EBNA2-}$ (*B*) virus. (*C*,*D*) Examples of EBNA2-dependent differentially expressed genes in the UCSC Genome Browser (hg19) showing up-regulated genes (*C*) and down-regulated genes(*D*) . (*E*,*F*) UCSC Genome Browser (hg19) depicting the EBV-dependent EBNA2-independent up-regulated genes (*E*) and down-regulated  genes (*F*).
