## Supplemental Figure S3 for "Epstein-Barr virus nuclear antigen 2 (EBNA2) extensively rewires the human chromatin landscape at autoimmune risk loci"

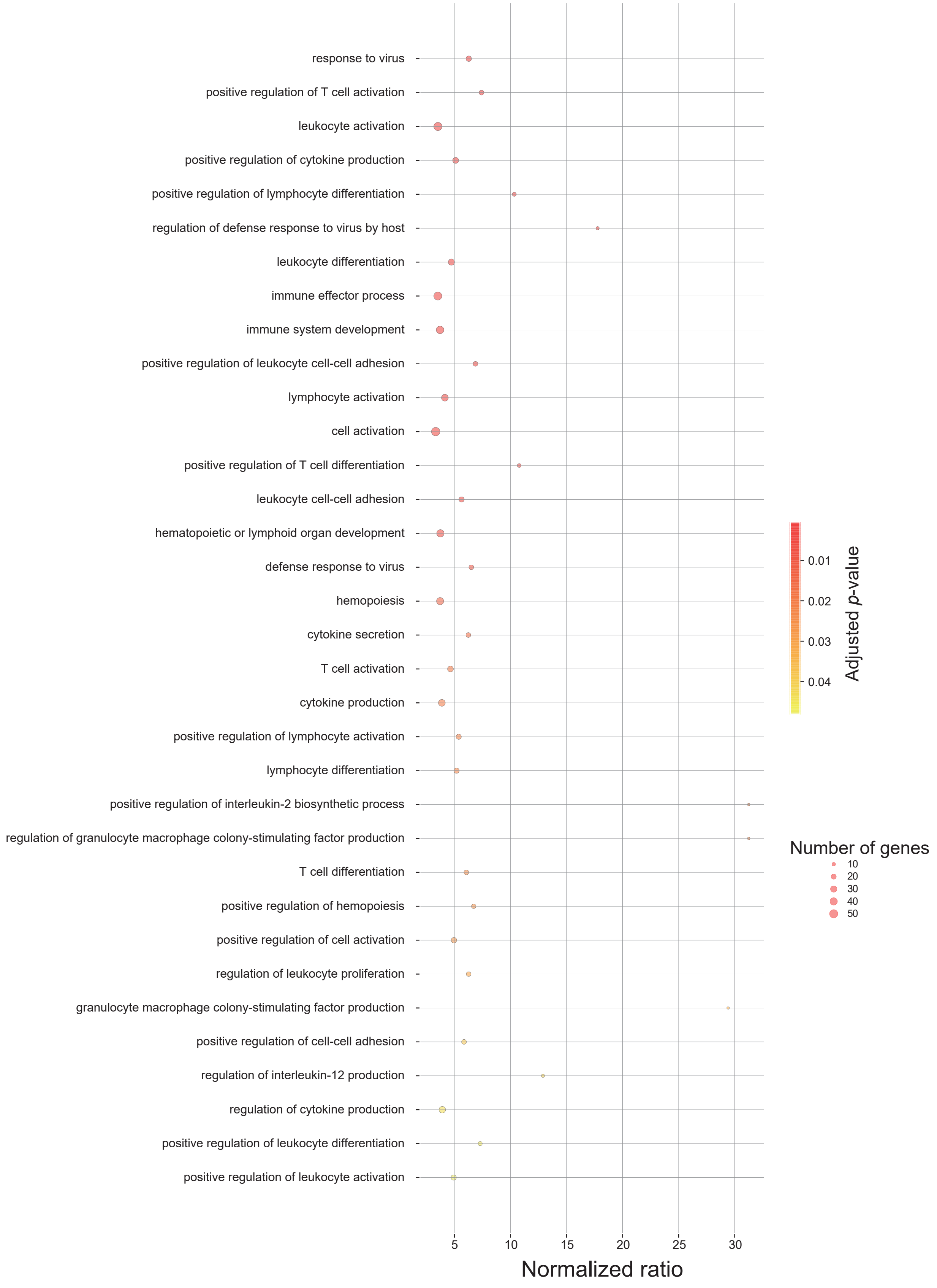


**Supplemental Figure 3**. Functional enrichment analysis for EBNA2-dependent differentially expressed genes. Gene set enrichment analyses of EBNA2-dependent differentially expressed genes for GO Biological Process. The X-axis indicates the normalized ratio of the number of input genes to the total number of annotated genes times 100. The Y-axis is sorted by significance (i.e., lower p-values are at the top of the plot).Analysis was performed using ToppFun from ToppGene Suite (https://toppgene.cchmc.org/enrichment.jsp). Bonferroni correction was used to adjust the *p*-values.
