## Supplemental Figure S4 for "Epstein-Barr virus nuclear antigen 2 (EBNA2) extensively rewires the human chromatin landscape at autoimmune risk loci"

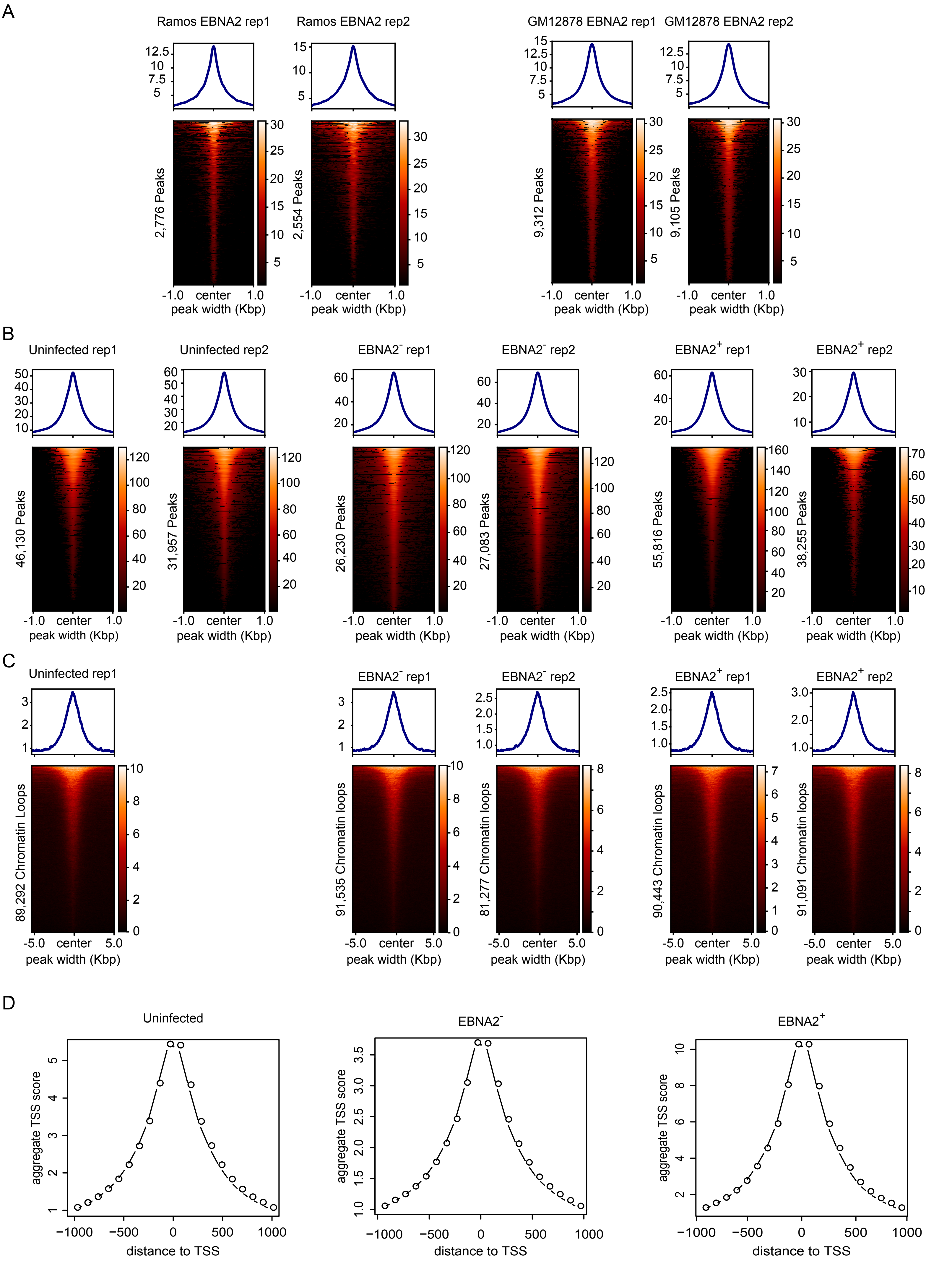


**Supplemental Figure S4. Data reproducibility and quality control**. Values in heatmaps indicate normalized read counts per genomic region and were generated using the computeMatrix tool in the deepTools package (Ramirez et al. 2016). Rows are matched between replicates (i.e., the same row indicates the same genomic locus between the replicates). Plots at the top indicate the average signal across the genomic loci. (A) Comparison of Ramos EBNA2 ChIP-seq replicates (left) and GM12878 EBNA2 ChIP-seq replicates (right). (B) Comparison of ATAC-seq replicates in uninfected (left), EBNA2- (center), and EBNA2+ (right) conditions. (C) Comparison of Hi-ChIP replicates in uninfected (left), EBNA2- (center), EBNA2+ (right) conditions. Only a single sample is available for the uninfected condition. (D) TSS enrichment score for the ATAC-seq samples, in uninfected (left), EBNA2- (center), and EBNA2+ (right) conditions. The TSS enrichment score was calculated using the ATACseqQC package (Ou et al. 2018).
