## Supplemental Figure S5 for "Epstein-Barr virus nuclear antigen 2 (EBNA2) extensively rewires the human chromatin landscape at autoimmune risk loci"

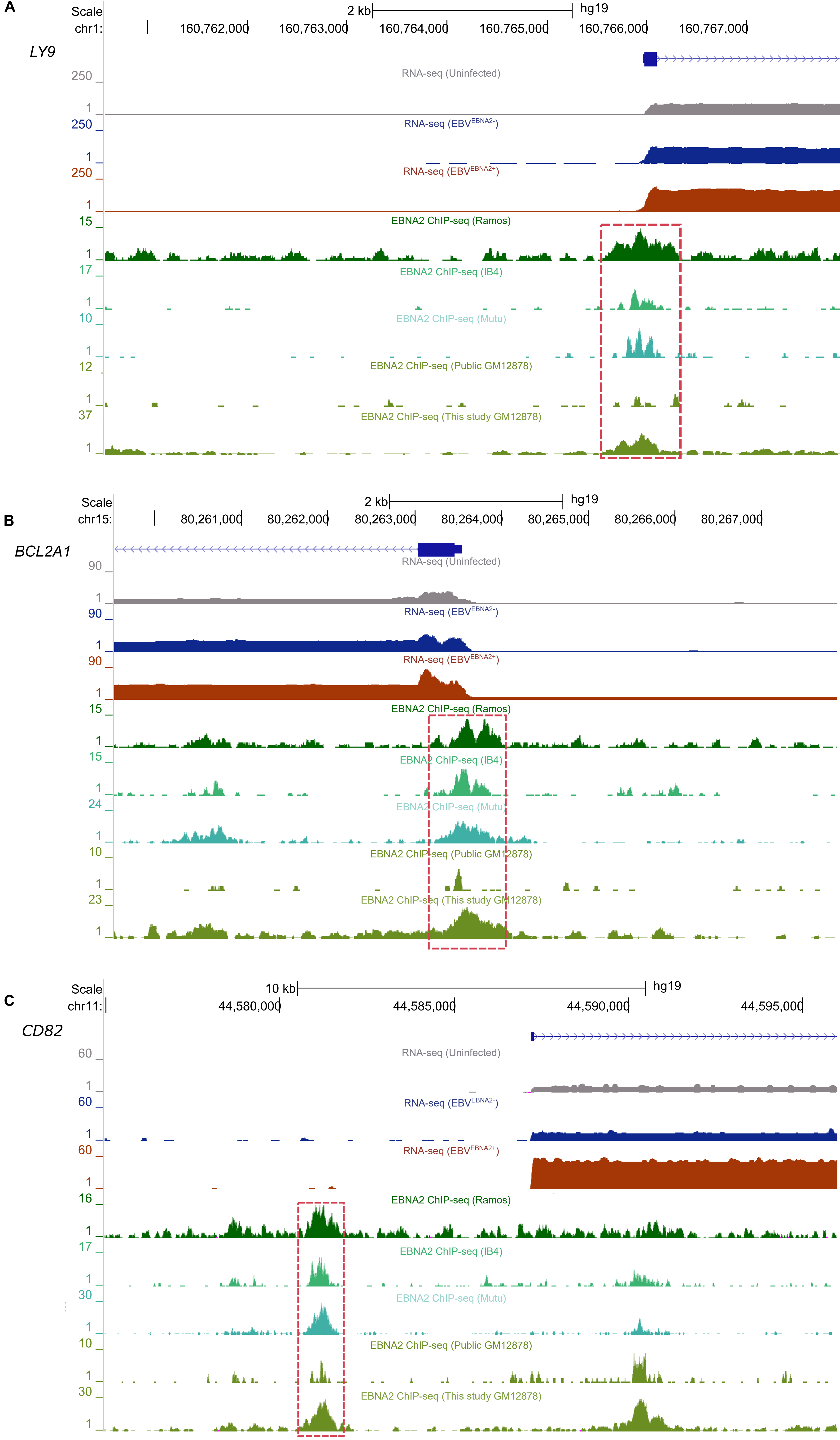


**Supplemental Figure 5.** EBNA2 binding at the promoter of EBNA2-dependent up-regulated genes (A-C). UCSC Genome Browser shots (hg19) for the LY9 (A), BCL21A (B), and CD82 (C) promoter regions. EBNA2 ChIP-seq peaks are boxed in green.
